# VeloTrace Reconciles Divergent Velocity and Trajectory in Single-cell Transcriptomics with Deep Neural ODE

**DOI:** 10.64898/2026.04.09.717458

**Authors:** Hui Cheng, Yunhao Qiao, Yufan Feng, Yuyang Wei, Jiachen Li, Jinpu Cai, Shuangjia Zheng, Siheng Chen, Guanbin Li, Benjamin D. Simons, Qiuyu Lian, Hongyi Xin

## Abstract

Cellular identity and fate transitions are governed by continuous molecular processes that form dynamic trajectories within a high-dimensional transcriptomic landscape. Existing methods attempt to model these dynamics from two complementary perspectives: trajectory inference and velocity modeling. Ideally, velocity and trajectory are dual aspects of transcriptomic dynamics where velocity is tangent to trajectory everywhere. This inherent connection between velocity and trajectory is currently absent in transcriptomic analysis. Splicing velocity are precision-limited to inadequately-sequenced genes, while trajectory inference prioritizes the modeling of global trends while omitting local dynamics. This divergence breaks the geometric continuity between local velocities and global trajectories, hindering the reliable interpretation of developmental dynamics. To reconcile trajectory inference and RNA velocity, we introduce VeloTrace, a framework that unifies them through Neural Ordinary Differential Equations (NeuralODEs). VeloTrace learns a continuous-time velocity field whose integral curves constitute the trajectory itself, while ensuring that velocities are tangent to integral paths everywhere. Leveraging a splicing quality score, VeloTrace incorporates high-quality splicing velocity as partial supervision for velocity orientation and grounding. During optimization, VeloTrace incorporates a Monte Carlo multi–time-frame supervision strategy to ensure coherence between local and global trajectorys and suppress sequencing-induced stochastic diffusion. Through refining the velocity field and cell-specific parameters for pseudo-time, expression, and velocity, VeloTrace reconstructs a smooth, local-and-global-coherent velocity-vector-guided flow in the transcriptomic latent space. This strategy ensures a complementary integration of velocity and trajectory, imputing the transcriptional kinetics for genes of insufficient strength, whose kinetics cannot be accurately portrayed by splicing velocity. In simulation benchmarks, VeloTrace captured the transcriptional dynamics of all expressed genes, even those with inadequate sequencing coverage, producing velocity directions that were most consistent with the true direction and everywhere tangential across the entire process, outperforming state-of-the-art methods, including scVelo, UniTVelo, VeloVI and scTour. VeloTrace uniquely reconciles RNA velocity and trajectory inference, creating a velocity field where each cell can infer past and future transitions from its current state. Moreover, VeloTrace extends reliable velocity estimation to a broader set of genes. When applied to mouse neural stem cell differentiation data, it successfully recovers dynamics of driver genes for two developmental lineages, including those with low expression, shedding light on their regulatory roles during differentiation. This unified framework lays the foundation for more accurate modeling of gene regulation and cell fate decisions in complex biological systems.

## Introduction

The transcriptome space is a mathematical construct that represents the gene expression profile of individual cells in a high-dimensional coordinate system, where each dimension corresponds to the abundance of a particular transcript. In single-cell RNA sequencing experiments, millions of cells can be sampled and positioned in this space, revealing substantial heterogeneity in gene expression across the cell population. Because gene expression is subject to continuous modulation through transcriptional regulation, RNA processing, and degradation, modeling cellular behaviors such as differentiation or responses to external perturbations requires viewing this space not as static, but as fundamentally dynamic. We adopt the perspective that the transcriptome landscape should be represented by a velocity field, where each cell is associated with both a coordinate, which denotes the current transcriptomic state of the cell, and a velocity vector that captures the underlying regulatory forces driving its transcriptomic reprogramming. Conceptually, cellular development trajectories can be represented as smooth flows guided by this velocity field, where cells progress along paths that are integrals of the velocity vectors (Fig. 1a).

**Figure 1:**
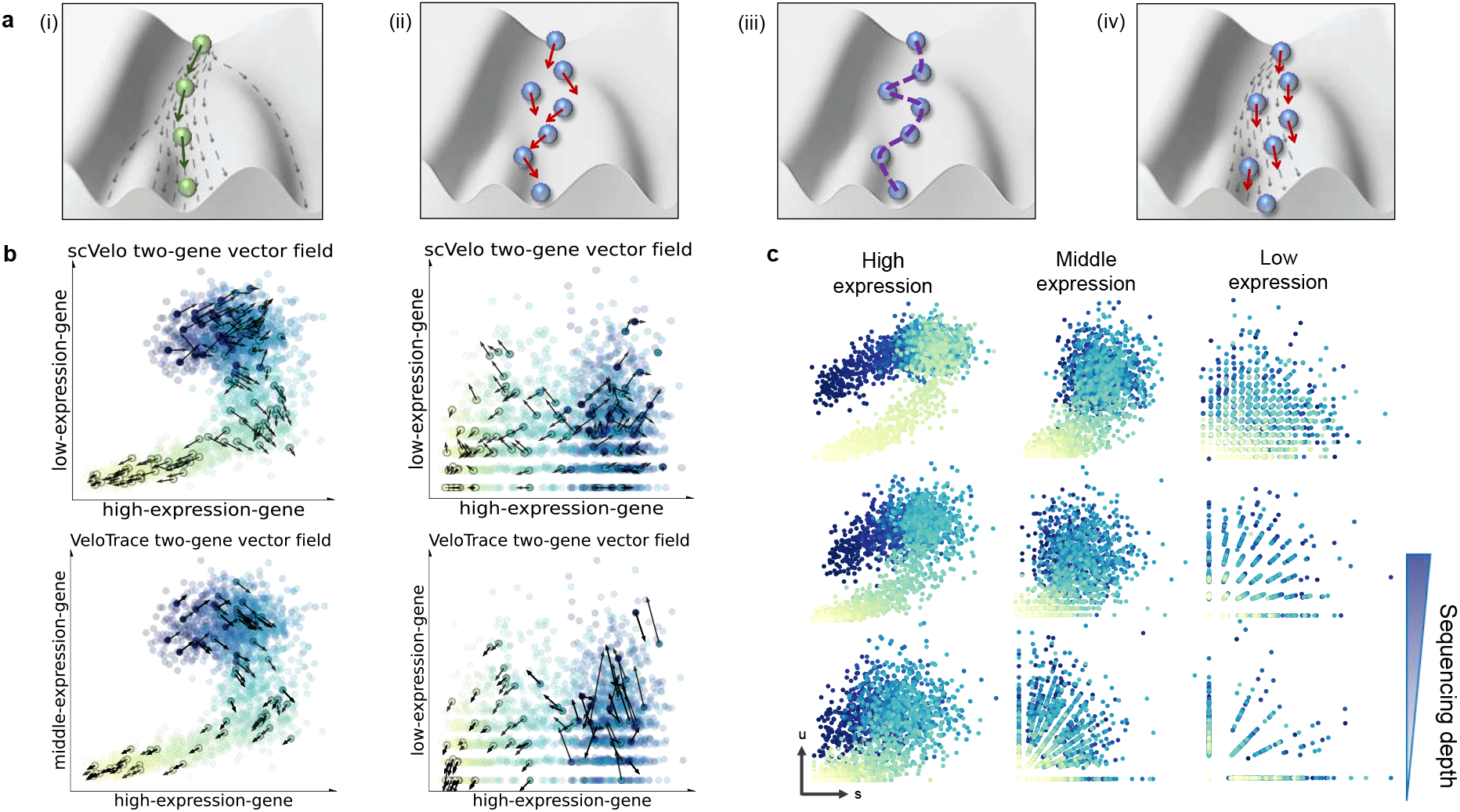
**a**, Conceptual illustration of the relationship between cell trajectories and velocity fields. Ideally, **i** the true developmental trajectory and its tangent velocity field are perfectly aligned. In real case, **ii** the velocity field inferred from scRNA-seq often exhibits local noise and direction inconsistency, and **iii** trajectory analysis connecting cell states with a smooth path in dimensional-reduced space does not encode local transcriptional directionality. **iv**, Constructing a velocity field that coherently aligns with the trajectory can integrate the strengths of both analyses. **b**, Demonstration of a two-gene example across different expression levels. Scatter plots of high-, intermediate-, and low-expression gene pairs show intuitive trajectory shapes in the 2D expression space, with inferred velocity vectors overlaid to illustrate how gene abundance affects velocity estimation. **c**, Spliced (s) and unspliced (u) counts for genes across expression levels (columns) and varying sequencing depths (rows). Higher expression and deeper sequencing produce clearer transcriptional kinetics, whereas low expression or down-sampling leads to increased sparsity and noisier phase portraits.

**Figure 2.**
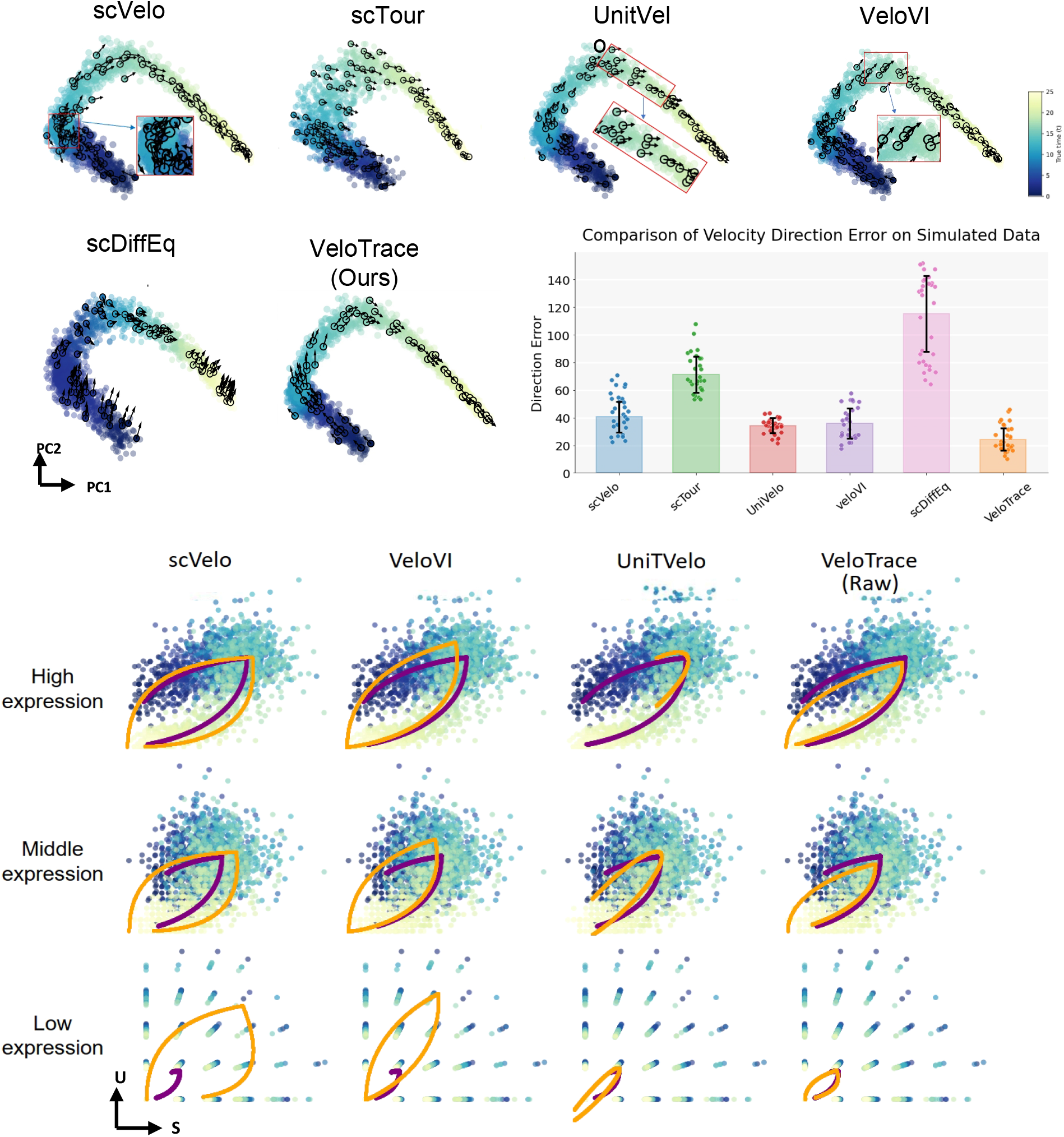
Benchmarking cell-wise and gene-wise velocity estimation and the coherence of the velocity field. **a**, Comparison of cell-wise velocity fields inferred by scVelo, VeloVI, UniTVelo, scTour, and VeloTrace. Top: Rose plots showing the angular error distributions between inferred and ground-truth velocity directions. Bottom: Velocity vectors projected into the principal-component (PC) space of gene-expression profiles to visualize local–global concordance with the underlying trajectory (for scTour, PCs of its latent embeddings are shown at right side, as velocities are defined in latent space). **b**, Fitted (orange) versus ground-truth (purple) phase portraits for representative high-, intermediate-, and low-expression genes, shown in spliced–unspliced (s–u) coordinate space. **c**, Gene-wise mean-squared error (MSE), calculated by comparing each cell’s spliced–unspliced position with its predicted position on the fitted phase portrait.

The first step towards modeling transcriptomic dynamics is to reconstruct the velocity field from high-dimensional single-cell RNA-seq data. This reconstruction must overcome several inherent challenges presented by the nature of scRNA-seq measurements. First, scRNA-seq provides only a static snapshot of a heterogeneous cell population at a single time point, yielding a cross-sectional distribution of cells across the transcriptomic manifold without direct temporal information about individual cellular trajectories. Inferring directional dynamics from such cross-sectional data requires strong assumptions about the temporal ordering and continuity of cell states along developmental or response pathways. Second, the scRNA-seq process is inherently stochastic, and the resulting data suffer from substantial technical noise including dropout events (false zeros due to low capture efficiency), variable sequencing depth across cells, and mean-variance instability where low-abundance transcripts experience disproportionately high sampling noise. Third, velocity field reconstruction must be reconciled with gene-specific splicing kinetics. Splicing dynamics, the kinetics of pre-mRNA processing and mature RNA production, represent an important layer of transcriptional regulation at the individual gene level. A coherent transcriptomic velocity field should therefore be consistent with and informed by the splicing dynamics of constituent genes, ensuring that inferred cellular velocities reflect both changes in steady-state mRNA levels and the underlying splicing-dependent transcription kinetics that govern their dynamics.

The coherence between transcriptome-level dynamics and gene-level splicing kinetics remains inadequately integrated in current single-cell analytical pipelines. Currently, these two aspects are modeled largely in isolation: trajectory analysis infers both a path connecting various cell states identified in the transcriptome landscape and a temporal ordering of cells along this path, based primarily on proximity relationships between cell states in the high-dimensional expression space [1, 2, 3]. In parallel, splicing dynamics, also termed RNA velocity, focuses on modeling transcription, splicing, and RNA degradation kinetics at the level of individual genes, capturing the intrinsic transcriptional rates that govern mRNA accumulation [4, 5, 6]. The kinetics estimated for individual genes are then aggregated cell-wise to derive a transcriptome-wide velocity vector for each cell in the landscape. Current practice attempts to reconcile these two perspectives by projecting gene-level splicing velocities onto the inferred trajectory, allowing RNA velocity estimates to establish directionality along the trajectory path and predict cell fate transitions.

Insufficient integration between trajectory inference and RNA velocity estimation creates latent conflicts between displacement and velocity in the transcriptome manifold. By definition, velocity represents the time derivative of position along a trajectory, and should therefore be tangent to the trajectory path everywhere. Currently, this tangency constraint is enforced in a post-hoc manner through projection of gene-level velocities onto an inferred trajectory, which masks underlying disagreements between the two frameworks. The discordance between independently inferred trajectories and RNA velocities has received attention in the field. A recent systematic analysis by Zheng et al. [7] evaluated the consistency of RNA velocity inference algorithms and found that velocity estimates are highly sensitive to sequencing noise and stochastic dropouts. Critically, this sensitivity often goes unrecognized because splicing-derived velocities are projected onto a pre-computed trajectory and are subsequently smoothed, creating an artificial appearance of coherent flow. Fig. 1b shows the raw RNA velocities projected to a 2-gene transcriptome sub-space. Without projection and smoothing, raw velocities struggle to maintain good alignment with the trajectory path, with rather chaotic and contradictory velocities in local pockets. More importantly, the velocity field deteriorates as we move from high-expression genes to low-expression genes, highlighting its sensitivity with respect to gene expression strength and sequencing depth.

Ordinary differential equations (ODEs) provide a powerful framework for modeling transcriptome dynamics by describing the rate of change of gene expression as a function of the cell’s current transcriptomic state and regulatory context. ODE natrually unify velocity and trajectory: velocity is mathematically defined as the time derivative of trajectory, and trajectory is the time integral of velocity. ODEs have been extensively applied across scientific disciplines, from fluid dynamics and electromagnetism to thermodynamics and chemical kinetics, where they serve as the foundational framework for understanding dynamic systems. In the context of single-cell transcriptomics, ODE-based approaches have emerged as powerful tools for capturing gene expression dynamics. Splicing-based velocity methods model the kinetics of transcription, splicing, and RNA degradation through coupled ODEs describing the dynamics of nascent and mature mRNA pools, providing insight into instantaneous transcriptional rates [4, 5]. Regulatory ODE models such as scKINETICS integrate gene regulatory networks with transcriptome dynamics, using linear ODEs constrained by TF-target interactions to model expression transitions along developmental trajectories [6]. Neural ODE approaches such as scTour employ deep learning-parameterized differential equations to capture complex, non-linear transcriptome dynamics and enable imputation of intermediate or missing cell states along continuous developmental paths [8]. Despite these successes, a critical limitation remains: current ODE-based methods lack principled mechanisms for enforcing coherence between transcriptome-level dynamics (gene expression changes) and gene-level splicing dynamics (transcription and degradation kinetics). Without appropriate regularization constraining the model, ODE frameworks are prone to overfit the data, yielding overly flexible and locally erratic trajectories where fine-grained transcriptional transitions are inconsistent with the coarse-grained developmental directions.

To address these challenges, we propose VeloTrace, a unified neural ODE framework that grounds velocity predictions in gene-level splicing kinetics while enforcing consistency between local transcriptional dynamics and global developmental structure. VeloTrace reconstructs a smooth, continuous-time velocity field over the transcriptome manifold such that transcriptomic state and velocity are jointly inferred through ODE integration, ensuring trajectories emerge naturally from the velocity field rather than being imposed post-hoc. To enforce coherence between transcriptome-level dynamics and gene-level splicing kinetics, VeloTrace introduces a Splicing Quality Score (SQS) that identifies high-confidence splicing velocity estimates for use as partial supervision in ODE-based transcriptional velocity inference. This principled integration of splicing information regularizes the ODE framework, preventing overfitting to noisy or contradictory signals. To resolve local-global inconsistency and avoid degenerate tumbling trajectories in local regions, VeloTrace employs a Monte Carlo sampling strategy where multiple segments of varying temporal lengths are sampled to constrain local velocity estimates. By randomly sampling multiple initial and terminal conditions across overlapping time intervals, VeloTrace mitigates stochastic artifacts arising from sequencing noise and prevents the ODE from overfitting to spurious or ambiguous cell orderings. Experiments on synthetic and real datasets demonstrate that VeloTrace accurately reconstructs transcriptional velocities across the full expression spectrum, capturing both robust high-abundance genes and volatile low-abundance transcripts. VeloTrace produces locally and globally smooth, internally consistent trajectories and yields biologically interpretable velocity vector fields in the transcriptome landscape. Application to cerebellar development data reveals lineage-specific driver genes controlling GABAergic and gliogenic differentiation, identifies critical early commitment state in GABAergic lineages, and uncovers proliferation signatures associated with progenitor maintenance. These results demonstrate that jointly modeling splicing kinetics and transcriptome dynamics substantially improves trajectory quality and biological interpretability in single-cell transcriptomic studies.

## Results

### Overview of VeloTrace

We developed VeloTrace, a framework that reconciles trajectory inference and RNA velocity by learning a continuous transcriptomic vector field with Neural Ordinary Differential Equations (Neural ODEs) directly in high-dimensional gene-expression space. Formally, the flow is defined as

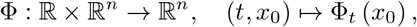

where cellular state transitions follow

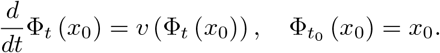

Under this formulation, cell trajectories arise as integral curves of a shared velocity field, allowing local transcriptional change and global developmental progression to be modeled within a single dynamical system.

To constrain the learned flow with biologically meaningful transcriptional regulation, we incorporated gene-specific splicing kinetics as supervision. For each gene, we fitted a kinetic model to unspliced and spliced counts and quantified its reliability using a splicing quality score (SQS), defined as the coefficient of determination between observed and predicted splicing dynamics across cells. Only high-confidence genes with sufficiently high SQS were retained to construct reference RNA-velocity vectors. We then enforced directional agreement between the Neural ODE field and these reference velocities using a cosine-alignment loss on the selected genes, such that the reconstructed flow remained consistent with reliable splicing-derived dynamics while avoiding noisy signals from poorly fitted genes. The model was optimized by combining this splicing-based regularization with the geometric reconstruction objective and the backward-consistency constraint, yielding a vector field that is both globally coherent and biologically directional.

### VeloTrace reconstructs conherent velocity field, extending reliable inference across gene expression levels

We began by assessing whether the velocity field inferred by VeloTrace accurately captures both local and – transcriptomic dynamics. We compared its performance on simulation datasets against three splicing-based methods (scVelo, VeloVI, and UniTVelo), the ODE-based method scTour and heuristic velocity estimates derived from trajectory and pseudotime. Because scTour learns velocity in an autoencoder-defined latent space, we decoded its latent-space velocities back to gene space to enable direct benchmarking against the ground truth.

Cell-wise velocity recovery is evaluated using the cosine distance between inferred and ground-truth velocity directions, with values converted into 0-180 degrees for visualization. To examine local–global concordance of the velocity field, we projected cells into a two-dimensional principal component (PC) space using their gene-expression profiles to visualize the overall trajectory, and overlaid the projected start and end points of the velocity vectors in the same space. Because PCA is a linear projection, it preserves local directional relationships more faithfully than nonlinear embeddings, enabling meaningful comparison between the projected velocity vectors and the underlying trajectory.

The benchmarking results demonstrate that VeloTrace provides the closest match to the groundtruth velocity field, and reconstructs a continuous and globally coherent flow along the trajectory. The angular error distributions in the top panel of Fig. 3a show the cell-wise discrepancies between the inferred and true velocities. The bottom panel illustrates the alignment between the overall trajectory and the inferred velocities. scVelo displays broad angular dispersion and produces locally inconsistent, chaotic vectors. VeloVI and UniTVelo show improved performance but still exhibit systematic angular shifts and region-specific misalignment between velocity and trajectory. scTour maintains tangency between velocity and trajectory in the latent space where it is trained (right side of its bottom panel), but once decoded to gene space, its velocities show wide angular deviations with many approaching 180 degree (top panel), indicating that the latent-space ODE dynamics do not translate to correct gene-expression dynamics or the true biological direction of transcriptional change In contrast, VeloTrace yields minimal angular error and produces smooth, directionally consistent vectors along the entire trajectory, faithfully capturing both local transitions and the overarching developmental flow.

**Figure 3.**
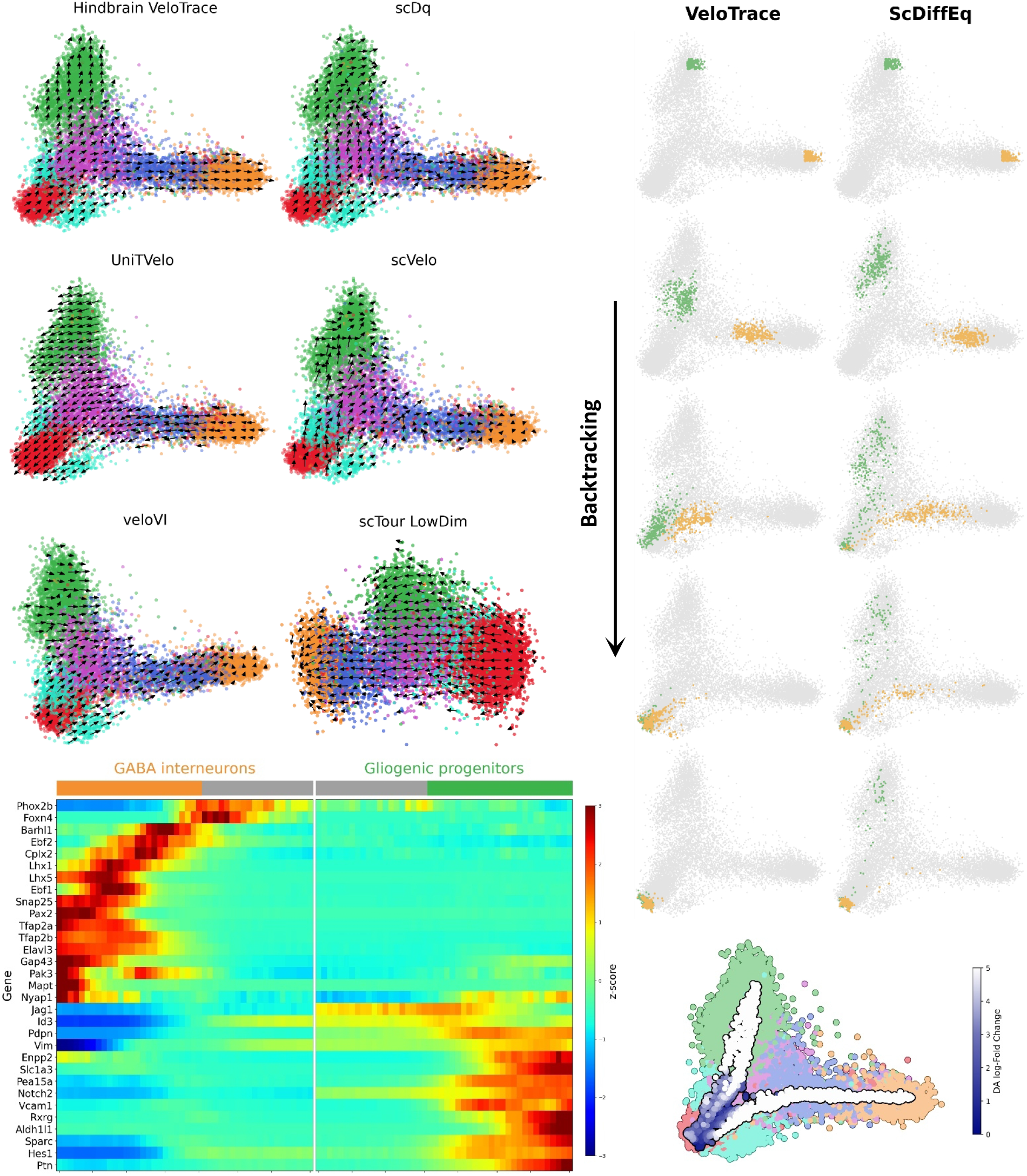
Application in a real scRNA-sea dataset of mouse cerebellar development. **a**, Prior knowledge of cell differentiation hierarchy and velocity vectors projected into the PC space (using gene-expression profiles for all methods except scTour, which uses PCs of its latent embeddings). **b**, Raw and VeloTrace-refined gene expression of Pax2 in two lineages (top: Glial, bottom: GABA). **c**, Dynamics-informative genes selected by Splicing Quality Score (SQS) in two lineages (top: Glial, bottom: GABA), and their corresponding raw and VeloTrace-refined gene expression patterns.

**Figure 4.**
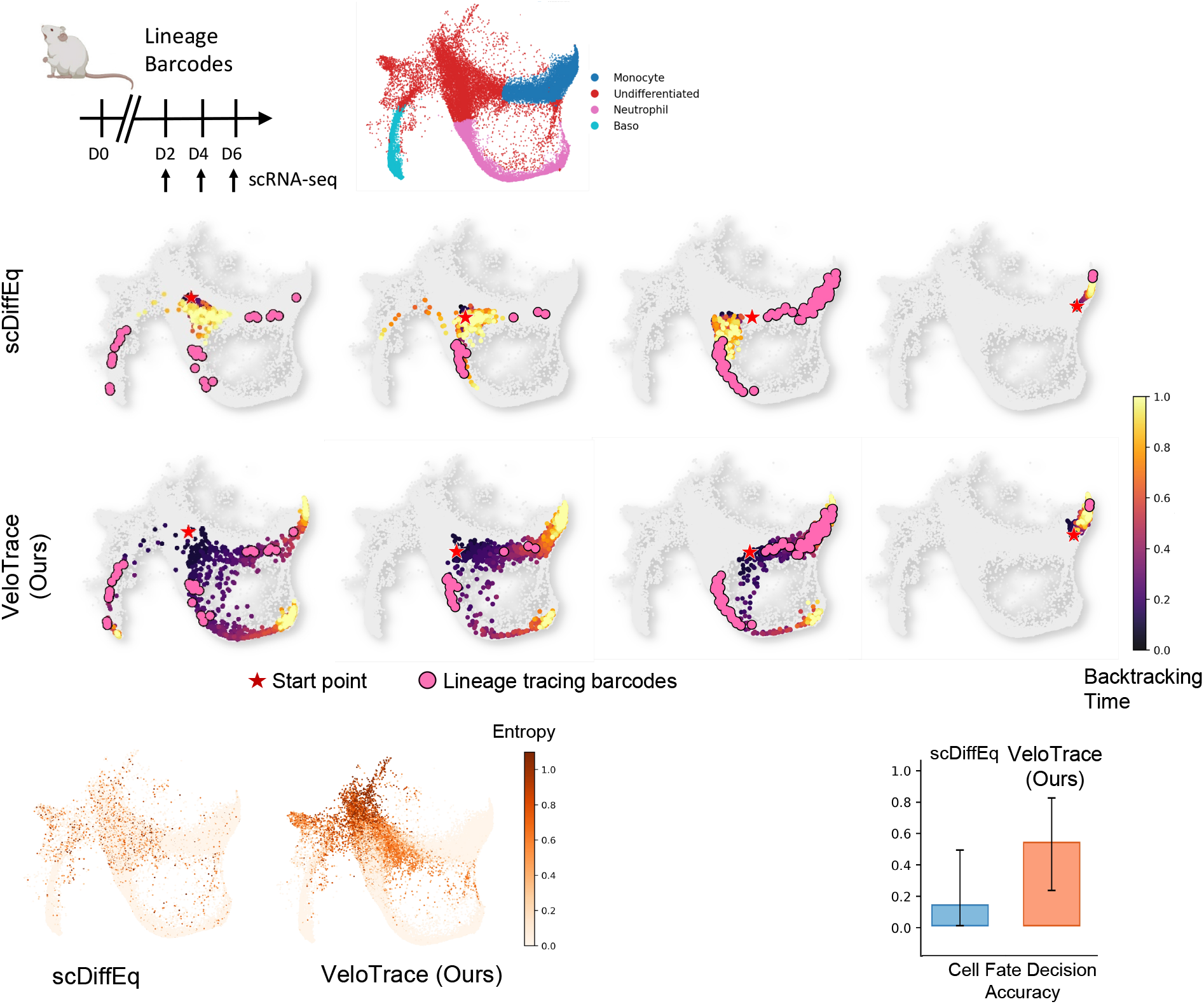
Application in a mouse hematopoietic development with lineage barcodes. **a**, Prior knowledge of cell differentiation hierarchy and velocity vectors projected into the PC space (using gene-expression profiles for all methods except scTour, which uses PCs of its latent embeddings).

In addition to velocity-specific methods, a straightforward and widely used approach is to derive velocity from trajectory inference and pseudotime. Here, Slingshot is used as a representative trajectory–pseudotime method due to its broad adoption. Conceptually, gene expression is modeled as a function of pseudotime, and changes in gene expression between adjacent pseudotime points are used to approximate velocity. However, because scRNA-seq data are highly noisy and sparse, cell-level estimates are extremely unstable, producing mosaic velocity vectors whose directions fluctuate frequently (Fig. 3b), indicating limited power for cell-wise velocity inference. Smoothing strategies, such as cluster-based averaging or fitting Generalized Additive Models (GAMs), are therefore required to recover the trends, but they introduce additional parameters that must be manually tuned. The resulting velocities are highly sensitive to these choices, and optimal parameter settings often vary across genes, making it nearly impossible to construct a robust, genome-wide velocity field using pseudotime-based approaches. Thus, trajectory/pseudotime-based velocity serves mainly as a qualitative descriptor under careful tuning, but does not support reliable quantitative velocity analysis.

In summary, VeloTrace most accurately recovers cell-wise velocities, and the vector field it constructs best preserves tangency to the trajectory and local–global coherence, demonstrating strong ability to capture transcriptional dynamics and infer biologically meaningful state transitions.

To evaluate gene-level velocity accuracy, we compared VeloTrace against scVelo, VeloVI, and UniTVelo– methods that directly support gene-level velocity estimation–using simulation data with known kinetic parameters and ground-truth phase portraits. In Fig. **??**a, the rows show fitted phase portraits produced by each method for representative high-, middle-, and low-expression genes. Across all expression levels, scVelo and VeloVI show varying degrees of deviation, particularly for low-expression genes, where noise is more prominent. UniTVelo shows smaller deviations from ground truth and behaves consistently across gene expression levels. VeloTrace refines gene-wise dynamics via time-guided Laplacian smoothing (see Methods) and significantly enhances gene-specific splicing trajectories by reducing noise and drawing cells closer to the ground-truth curve, even for middle- and low-expression genes.

To quantitatively summarize the performance, we computed the mean-squared error (MSE) between cellular unspliced and spliced counts (*u*_*i*_, *s*_*i*_) and the predicted values on the fitted phase portrait 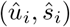 at assigned kinetic time points: 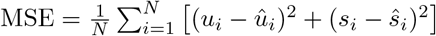, where *N* is the number of cells. The results in Fig. **??**b) show that across high-, intermediate-, and low-expression gene groups, scVelo and VeloVI showed substantially higher and more variable errors, demonstrating limited robustness in recovering gene-specific velocity dynamics. UniTVelo and the raw-count version of VeloTrace achieve MSE values comparable to those obtained under ground-truth kinetics (where the MSE reflects only measurement noise). The refined VeloTrace further reduces MSE across all expression levels and shows a pronounced improvement for low-expression genes, where noise is greatest. These results demonstrate that VeloTrace robustly recovers gene-specific velocity dynamics regardless of gene abundance, and that its refinement based on the constructed vector field provides significant enhancement of gene-dynamic patterns.

### Retrospective cell fate prediction identifies early commitment states of cerebellar development

To further evaluate the capability of VeloTrace in resolving cell fate, we applied it to a bifurcating dataset of mouse cerebellum cell development. In this study, VeloTrace, again, effectively constructs the velocity field and recovers the kinetics and transcriptome of cells. Compared to existing methods, VeloTrace out-performs in capturing dynamic cellular processes across diverse cell types, providing enhanced accuracy in resolving cellular transitions (Fig. **??**a). In contrast, scVelo displays local inconsistencies within the ventricular zone (VZ) progenitors, with inferred trajectories deviating from the expected directionality in certain regions. The velocity plot generated by VeloVI contradict the anticipated differentiation flow of gliogenic progenitors. ScTour fails to properly distinguish between gliogenic and GABA interneuron lineages, mistakenly classifying gliogenic progenitors as a source for GABA interneurons. Meanwhile, UniTvelo, despite being locally consistent, shows a complete reversal in the directionality of the velocity, diverging from the true trajectory.

In the gliogenic lineage, the refinement of gene dynamics yields similar improvements. Genes like Ptprz1, which regulates oligodendrocyte differentiation and glial development signaling and Slc1a3, an astrocyte marker and regulator of glutamate homeostasis, shows a significant expression pattern observed across progenitor stages follow their expected dynamics [9, 10].

In terms of proliferation and differentiation, several key genes are revealed through imputation. Rpl4, a protein synthesis gene, and Rrm2, involved in DNA synthesis and associated with glial tumors (gliomas) and poor prognosis, show reduced expression during gliogenic differentiation, highlighting their roles in controlling proliferation during neural stem cell differentiation. Set expression decreases during differentiation, marking a reduction in proliferative and undifferentiated states. Meanwhile, Sox6, which maintains a proliferative or immature state, and Ddah1, supporting glial progenitor differentiation, are upregulated in gliogenic progenitors. These two gene sets act antagonistically, regulating proliferation during differentiation.

In summary, VeloTrace outperforms in resolving cell fate by effectively constructing velocity fields and recovering gene kinetics, aligning gene expression with expected patterns across diverse cell types.

### VeloTrace predicts cell fate and reveals the thermodynamic landscape along developmental trajectories

We next evaluated VeloTrace on the hematopoietic lineage-tracing dataset, in which expressed barcodes link early transcriptional states to later fate outcomes measured by scRNA-seq across days 2, 4 and 6. To enable robust clonal comparison, we focused on three well-sampled branches—monocyte, neutrophil and basophil.

Against this benchmark, VeloTrace showed higher cell fate probability concordance, measured as the cosine similarity between predicted fate distributions and barcode-derived lineage outcomes. For early undifferentiated cells, scDiffEq displayed weak drift near the progenitor hub, with trajectories remaining trapped around the starting region rather than traversing the differentiation manifold. By contrast, VeloTrace produced broader, directed paths that aligned more closely with barcode-supported ancestral distributions. This advantage is more apparent for transitional cells downstream of the undifferentiated compartment. scDiffEq frequently produced fate assignments that were only partially consistent with the barcode-supported outcomes, including false-positive basophil predictions and underestimation of monocyte-directed potential in cells with mixed lineage bias. By contrast, VeloTrace generated predictions that more closely followed the fate composition of each starting cell, preserving shared monocyte–neutrophil potential where fate choice remained unresolved.

## Disscussion

This study introduces VeloTrace, a unified framework that integrates RNA velocity and trajectory inference to provide a continuous, coherent representation of transcriptomic dynamics. By leveraging Neural Ordinary Differential Equations (NeuralODEs), VeloTrace ensures that the velocity field is tangential to the trajectory at every point, addressing a key gap in existing methods. While traditional trajectory inference captures global trends, it overlooks local dynamics, and splicing velocity is often limited by sequencing depth, especially for low-expression genes. VeloTrace overcomes these challenges by incorporating high-quality splicing velocity as partial supervision and utilizing Monte Carlo multi-time-frame supervision to maintain consistency between local and global dynamics, while mitigating sequencing-induced noise.

In simulation benchmarks, VeloTrace demonstrates superior performance in recovering local- and global-coherent transcriptional dynamics at both gene- and cell-wise levels. It consistently outperformed state-of-the-art methods, including scVelo, scTour, and DeepVelo, in terms of accuracy and consistency with true transcriptional trajectories. The framework’s ability to recover accurate velocity directions and gene kinetics, even for genes with insufficient sequencing coverage, represents a significant advancement over existing approaches.

When applied to mouse neural stem cell differentiation, VeloTrace successfully refines dynamics patterns of key driver genes for two developmental lineages, highlighting its potential for uncovering regulatory roles during differentiation. This ability to reliably estimate velocity and trajectory across the transcriptome enhances our understanding of cellular fate transitions and gene regulation.

Despite its strengths, VeloTrace’s network architecture lacks sufficient interpretability, which limits its ability to fully integrate gene regulation and signaling pathway information. Future improvements could focus on incorporating these layers of biological context to enhance the framework’s explanatory power. Nonetheless, VeloTrace represents a significant step forward in transcriptomic modeling, offering a more comprehensive framework for understanding cellular dynamics, gene regulation, and cell fate decisions in complex biological systems.

## Methods

### Formalizing the Transcriptomic Velocity Field

In dynamical scenarios, the evolution of a state *x*(*t*) over time is often described by a vector field *v* : ℝ^*n*^→ℝ^*n*^, where each point *x* in the system’s state space is associated with a velocity vector *v*(*x*), which determines the direction and rate of change of the state at that point. The trajectory of the system can be defined as a flow Φ_*t*∈*R*+_ (*x*_0_) governed by this velocity field, which maps the initial state *x*_0_ to its position for a duration *t*.

In the context of transcriptomic dynamics, we extend this framework to model the evolution of cellular gene expression as a dynamical system. Here, the vector field **v**(**x**) represents the instantaneous transcriptional changes at state **x**, describing how cells progress through the transcriptomic landscape. The flow of this field, Φ_*t*_ (*x*_0_), captures the trajectory of a cell’s molecular state over time. Just as in general dynamical systems, this flow describes the progression of cells in gene expression space, where gene-level velocity fields govern the dynamics of transcriptional regulation.

We propose a high-dimensional velocity field model to describe transcriptomic dynamics, where the velocity field, *v* : ℝ^*n*^ →ℝ^*n*^, which assigns to each point *x* a velocity vector **v**(**x**), describing the instantaneous rate and direction of change at that location. The motion of a cell under this field defines a trajectory Φ_*t>*0_ (*x*_0_), which represents the position of the cell after evolving for a duration *t* from its initial state *x*_0_. Formally, the flow is a mapping

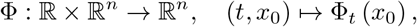

satisfying the ordinary differential equation (ODE):

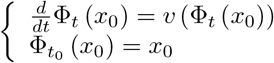

At each time *t*, the cell’s velocity is determined by the vector field at its current position. The set 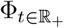 thus forms a flow describing how points move continuously through the velocity field. This high dimensional coherent transcriptomic velocity field should be smooth and consistent with gene-level velocity field.

However, in single-cell RNA sequencing, we are only able to observe discrete snapshots of individual cells, *x*_*i*_, rather than continuous trajectories *x*(*t*). To model the underlying complex and nonlinear dynamics, we leverage Neural Ordinary Differential Equations (Neural ODEs). Neural ODEs provide a framework to learn a continuous velocity field from discrete observations, enabling the reconstruction of smooth, continuous trajectories for cellular states. This approach allows us to define a globally consistent vector field that aligns the estimated gene kinetics with the cell-wise velocity field, offering a continuous and coherent description of the transcriptomic trajectory over time.

### Transcriptomic Velocity Field Reconstruction Grounded by High Quality Splicing Velocity

To reconstruct this implicit dynamical flow from static single-cell data, we modeled the transcriptomic state transitions as a Neural Ordinary Differential Equation (NeuralODE),

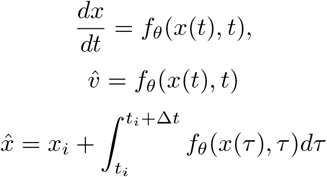

where *f*_*θ*_ denotes a neural network parameterizing the local velocity field on the transcriptomic manifold. In principle, the temporal evolution from state *x*_*i*_ to *x*_*i*+1_ follows the flow map

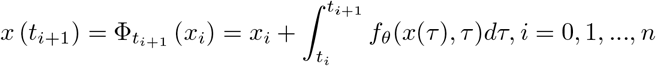

The geometric supervision loss is defined as:

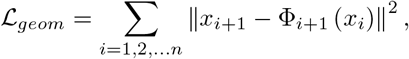

which constrains the NeuralODE to learn a globally consistent flow across observed transcriptomic states.

In biological context, velocity field reconstruction should also be consistent with gene-specific splicing kinetics, an essential layer of transcriptional regulation that governs the production of mature mRNA from pre-mRNA. To ensure that the reconstructed flow aligns with biologically meaningful transcriptional regulation, we incorporate high-confidence splicing kinetics as a weak supervision signal using a splicing quality score (SQS). For each gene *g*, a kinetic model is fitted to its unspliced (*u*_*g*_) and spliced (*s*_*g*_) mRNA counts. Over the full set of inferred times 𝒯= {*t*_1_, *t*_2_, …, *t*_*C*_}, the coefficient of determination for gene *g* is defined as:

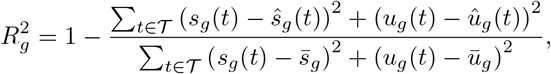

where *s*_*g,c*_ is the observed spliced count of gene *g* in cell *c, ŝ* _*g,c*_ is the predicted spliced count from the scVelo kinetic model, 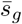 is the mean of observed spliced counts across all cells for gene *g*, and *C* is the total number of cells. A higher 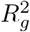 value indicates that the splicing kinetic model better explains the observed splicing dynamics for that gene.

Genes with poor fits 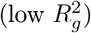 yield noisy velocity estimates and are excluded. We define a threshold *R*^2^ and the high-quality gene set as:

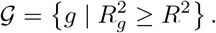

Using only this reliable gene set, reference velocity vectors 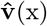 are constructed. To enforce directional agreement between predicted and reference velocities, we define velocity alignment loss using the negative cosine similarity:

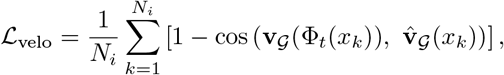

This regularization encourages the reconstructed vector field to follow biologically plausible directions guided by splicing kinetics.

The internal ODE integration is solved using an adaptive Runge–Kutta method. Model parameters are optimized using stochastic gradient descent over the combined objective:

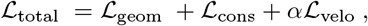

where *α* controls the trade-off between displacement fidelity and splicing-based alignment, which yields a vector field that is both geometrically coherent and biologically directional.

### Achieving Transcriptomic Coherency through Monte-Carlo Flow Sampling with Multi-Time-Lag

#### Monte-Carlo flow sampling for vector field estimation

Since single-cell measurements provide only instantaneous snapshots rather than continuous trajectories, we approximate the true flow integral using Monte Carlo sampling. Specifically, latent temporal coordinates are first estimated using a pseudotime inference method, providing a coarse ordering of cells along the developmental continuum. At each training iteration, starting states *x*_*i*_ and corresponding temporal displacements {∆*t*} _*i*_ are randomly sampled from the observed data distribution. For each temporal interval ∆*t* and associated cell pairs {(*x*_*i*_, *x*_*j*_)}were stochastically drawn to construct a surrogate supervision signal

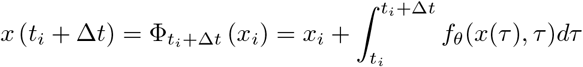

and the loss function is

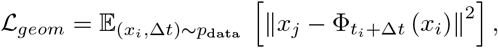

where 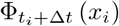 represents the NeuralODE-predicted flow integrated from *x*_*i*_ over ∆*t* to *x*_*j*_.

This stochastic formulation allows the model to approximate the true expectation over all possible start points and temporal displacements, thereby learning the average geometric flow consistent with the observed population distribution.

## Notes

### Competing Interest Statement

The authors have declared no competing interest.

